# Beyond the forest-grassland dichotomy: the gradient-like organization of habitats in forest-steppes

**DOI:** 10.1101/2020.01.12.903344

**Authors:** László Erdős, Péter Török, Katalin Szitár, Zoltán Bátori, Csaba Tölgyesi, Péter János Kiss, Ákos Bede-Fazekas, György Kröel-Dulay

## Abstract

Featuring a transitional zone between closed forests and treeless steppes, forest-steppes cover vast areas and have outstanding conservation importance. The components of this mosaic ecosystem can conveniently be classified into two basic types, forests and grasslands. However, this dichotomic classification may not fit reality as habitat organization can be much more complex. In this study, our aim was to find out if the main habitat types can be grouped into two distinct habitat categories (which would support the dichotomic description), or a different paradigm better fits this complex ecosystem. We selected six main habitats of sandy forest-steppes, and, using 176 relevés, we compared their vegetation based on species composition (NMDS ordination, number of common species of the studied habitats), relative ecological indicator values (mean indicators for temperature, soil moisture, and light availability), and functional species groups (life-form categories, geoelement types, and phytosociological preference groups). According to the species composition, we found a well-defined gradient, with the following habitat order: large forest patches – medium forest patches – small forest patches – north-facing edges – south-facing edges – grasslands. A considerable number of species were shared among all habitats, while the number of species restricted to certain habitat types was also numerous, especially for north-facing edges. The total (i.e., pooled) number of species peaked near the middle of the gradient, in north-facing edges. The relative ecological indicator values and functional species groups showed mostly gradual changes from the large forest patches to the grasslands. Our results indicate that the widely used dichotomic categorization of forest-steppe habitats into forest and grassland patches is too simplistic, potentially resulting in a considerable loss of information. We suggest that forest-steppe vegetation better fits the gradient-based paradigm of landscape structure, which is able to reflect continuous variations.

## 1 Introduction

Ecosystems where tree-dominated and grass-dominated patches form a mosaic (e.g., savannas, wood pastures and forest-steppes) cover a substantial proportion of Earth’s terrestrial surface (House et al., 2003), and their dynamics (e.g. Innes et al., 2013), biodiversity patterns (e.g. Erdős et al., 2018a, b), and conservation importance (e.g. Bergmeier et al., 2010; Prevedello et al., 2018) are in the focus of ecological studies. The components of such systems can conveniently be classified into two basic types, forests and grasslands, which differ substantially in several biotic (e.g., species composition and leaf area) and abiotic (e.g., solar radiation and soil moisture) parameters (Breshears, 2006). The presence of structurally dissimilar patches increases spatial heterogeneity and contributes to the maintenance of species diversity, ecosystem services, and ecological stability (Manning et al., 2009; Santana et al., 2017; Tölgyesi et al., 2018).

Forest-steppes form the contact zone between closed-canopy temperate forests and treeless steppes and provide a textbook example of forest-grassland mosaics (Erdős et al., 2018a). In these areas, alternating forest and grassland patches are an inherent feature of the ecosystem (Erdős et al., 2014; Hais et al., 2016; Bátori et al., 2018; Lashchinskiy et al., 2017). It has been highlighted that such complex ecosystems cannot be understood by simply studying the forest and grassland components independently (House et al., 2003; Erdős et al., 2018b). Instead, a holistic approach with an integrated view of the whole mosaic is needed, as these components are ecologically interrelated in several ways. For example, some animal species need both components for their full life cycles, while some plant species may switch preferences between the components in years with different weather patterns (Bartha et al., 2008; Luza et al., 2014).

Forests and grasslands have distinct environmental, structural and compositional characteristics, and the interactions of these characteristics result in the emergence of specific edge communities (Erdős et al., 2013, 2014). Edges are major habitats for tree recruitment; thus, they play an important role in forest-steppe dynamics (Erdős et al., 2015). These findings support the notion that, in addition to forest and grassland components, forest edges should also be recognized as important habitats in these mosaic ecosystems.

It is well known that treeless areas exert a considerable influence on the microclimatic parameters of the peripheral areas of forests, which may affect the whole area of small forest patches (Schmidt et al., 2017). Thus, it can be assumed that different-sized forest patches differ considerably regarding both structure and species composition.

As edges are transitional habitats between the forest and the grassland components, and small forest patches may in some characteristics be transitional between larger forest patches and grasslands, we hypothesize that the habitats of forest-steppe mosaics can be arranged along a gradient. However, this phenomenon in forest-steppes has so far received little scientific attention (Erdős et al., 2018b).

In this study, we evaluated six habitat types of sandy forest-steppes: large forest patches, medium forest patches, small forest patches, north-facing forest edges, south-facing forest edges, and grasslands. Our question was whether, based on species composition, functional species groups, and ecological indicator values, these six habitat types can be grouped into two distinct habitat categories (which would support the dichotomic description), or a different paradigm better fits this complex system.

## 2 Materials and Methods

### 2.1 Study area

We performed our study in the Kiskunság Sand Ridge (Central Hungary), near the westernmost extensions of the Eurasian forest-steppe belt. We selected six study sites where forest-grassland mosaics have been preserved in near-natural conditions: Csévharaszt (N 47°17’, E 19°24’), Tatárszentgyörgy (N 47°02’, E 19°22’), Fülöpháza (N 46°52’, E 19°25’), Bócsa (N 46°41’, E 19°27’), Tázlár (N 46°30’, E 19°30’), and Négyestelep (N 46°17’, E 19°35’). Each site is characterized by stabilized calcareous sand dunes of aeolian origin. The elevations of the sites vary between 105 and 140 m asl. The mean annual temperature is 10.0-10.7 °C, and the mean annual precipitation is 520-580 mm (Dövényi, 2010). Soils are humus-poor sandy soils with low water retention capacities (Várallyay, 1993).

The natural vegetation of the study sites represents a mosaic of forest and grassland patches. The grassland component of the vegetation mosaic is mainly formed by open perennial sand grasslands that are dominated by *Festuca vaginata*, *Stipa borysthenica*, and *S. capillata*. Other common species include *Alkanna tinctoria*, *Dianthus serotinus*, *Euphorbia seguieriana*, *Fumana procumbens*, *Koeleria glauca*, and *Potentilla arenaria*. The forest component is represented by differently sized patches of juniper-poplar stands. The canopy layer is 15-20 m high and is co-dominated by *Populus alba* and *P. × canescens*. The canopy is open, and its cover typically varies between 40 and 70%. The height of the shrub layer is 1-3 m, and its cover usually ranges from 20 to 80%. The most common shrubs are *Crataegus monogyna*, *Juniperus communis*, *Ligustrum vulgare*, and *Rhamnus cathartica*. The herb layer is sparse (10-40%) and is composed of species such as *Anthriscus cerefolium*, *Carex flacca*, *C. liparicarpos*, *Pimpinella saxifraga*, *Polygonatum odoratum*, and *Stellaria media*, as well as numerous tree and shrub seedlings. The edges are rather narrow, usually with extensive cover of shrubs (mainly *Crataegus monogyna* and *Juniperus communis*) and herbs (e.g., *Calamagrostis epigeios*, *Poa angustifolia*, *Teucrium chamaedrys*). The names of the plant species are according to Király (2009).

All study sites are under legal protection. Their current mosaic patterns are a result of the semiarid climate complemented by the extreme soil conditions. Evidence indicates that the spatial arrangement of the forest and grassland patches is stable, and the existence of the grassland component does not depend on grazing, fire, or other forms of disturbances (Fekete, 1992; Erdős et al., 2015). Due to the legal protection, anthropogenic disturbances in the study sites are minimal (a low level of non-destructive research and strictly regulated tourism). Natural disturbances include the effects of grazers and browsers (*Capreolus capreolus*, *Cervus elaphus*, *Dama dama*, and *Lepus europaeus*) as well as the activity of burrowing animals (*Meles meles*, *Talpa europaea*, and *Vulpes vulpes*), but their influence on the forest-grassland balance is presumably negligible. During the last decades, wildfire occurred in only one of the study sites but areas affected by the fire event were not sampled during our study.

### 2.2 Vegetation sampling

Within each site, we distinguished six habitat types: large forest patches (> 0.5 ha), medium forest patches (0.2-0.4 ha), small forest patches (< 0.1 ha), north-facing forest edges, south-facing forest edges, and grasslands. Within each habitat, 25-m2 plots were established. This size is small enough to sample even the smallest forest patches but large enough for a standard coenological relevé. The plot shape was 5 m × 5 m in the forest patches and grasslands, while we used 2 m × 12.5 m plots at the edges to ensure that the plots did not extend into forest or grassland interiors. Previous studies suggested that plot shape does not have distorting effects on the results at this scale (Keeley and Fotheringham, 2005; Bátori et al., 2018). An edge was defined as the zone outside of the outermost tree trunks but still under the canopy. For edge plots, only forest patches > 0.2 ha were taken into consideration.

We sampled a total of 176 plots: 27 plots in large forest patches, 29 plots in medium forest patches, and 30 plots each in small forest patches, north-facing edges, south-facing edges, and grasslands. Within each plot, the percentage cover of all vascular plant species of each vegetation layer was estimated visually in April and July 2016. For each species, the largest cover value was used for subsequent data analyses. Percentage cover values for all species recorded in the 176 study plots are given in Supplementary Data A1.

### 2.3 Data analyses

We performed Non-metric Multidimensional Scaling (NMDS) to two dimensions, a robust unconstrained ordination method widely applied in community ecology (Minchin, 1987), based on the presence-absence data of the species, using the Sørensen–Dice index. For the analysis, the ‘vegan’ R package (Oksanen et al., 2018) was used. Several (minimum 500, maximum 5000) NMDSs were run from random starts to facilitate convergence to a non-local optimum. The result was visualized using centering, half-change scaling and rotation to the axes of Principal Components Analysis (PCA), hence the variance of the points is maximized on the first dimension.

To visualize the overlaps of the six studied habitats and assess their distinctness, we applied the technique of Lex et al. (2014). This method shows the number of species restricted to certain habitat types (i.e., habitat distinctness) as well as the number of species present in two or more habitats (i.e., habitat overlaps). Calculations were performed in MS Excel, and the graphs were prepared with Adobe Photoshop 7.0.

We calculated the mean ecological indicator values for temperature, soil moisture, and light availability for each plot. We used the indicator values of Borhidi (1995), which are based on the values of Ellenberg (1992) but extended for the Carpathian Basin. As proven by numerous field measurements, ecological indicator values are able to provide reliable estimates of site conditions (e.g., Schaffers and Sýkora, 2000; Dzwonko, 2001; Tölgyesi et al., 2014). Although the use of mean indicator values is often criticized, it has been shown that they perform well and have a solid theoretical basis (ter Braak and Gremmen, 1987; Diekmann, 2003). Ecological indicator values provide important information as they integrate fluctuating values over time (rather than providing data for a very short period) and reflect site conditions in relation to species’ requirements (rather than the mere absolute values of environmental parameters) (Zonneveld, 1983; Bartha, 2002; Diekmann, 2003).

Linear mixed effects models were used to test for the effects of habitat type on the mean indicator values of temperature, soil moisture and light availability. We used site as a random effect in the analyses. The statistical tests were implemented using the nlme R package (Pinheiro et al., 2013). The fulfillment of the normality and homoscedasticity assumptions of the models were checked by visual assessments of diagnostic plots. As model residuals showed heterogeneity of variances, we used a variance structure that allowed for different residual spreads for each level of explanatory variable according to Zuur et al. (2009). Tukey’s HSD tests were implemented for post hoc pairwise comparisons of the habitat categories using the multcomp R package (Hothorn et al., 2008).

Species were classified into life-form categories, geoelement types, and phytosociological preference groups. The categorizations were based on Horváth et al. (1995) and Borhidi (1995). Geoelement types reflect the global distribution of species, while phytosociological preference groups describe the regional preferences of species to certain plant communities. For phytosociological preferences, only the native species were considered, as non-natives tend to have indefinite preferences. The frequency distributions were calculated for all three categorizations for each habitat and were compared using Pearson’s chi-squared test.

All statistical analyses were performed in the R statistical environment ver. 2.15.2 (R Development Core Team, 2010).

## 3 Results

In the 176 plots, a total of 232 vascular plant species were found. The NMDS converged after 2267 tries and 90 iterations, achieving a stress value of 0.1784. The ordination revealed that the habitats were aligned in the ordination space following the sequence large forest patches – medium forest patches – small forest patches – north-facing edges – south-facing edges – grasslands (Figure 1). The species turnover was mostly gradual. Although the grassland habitat formed a relatively distinct group, its species composition showed some overlap with that of the south-facing edges.

**Figure 1.**
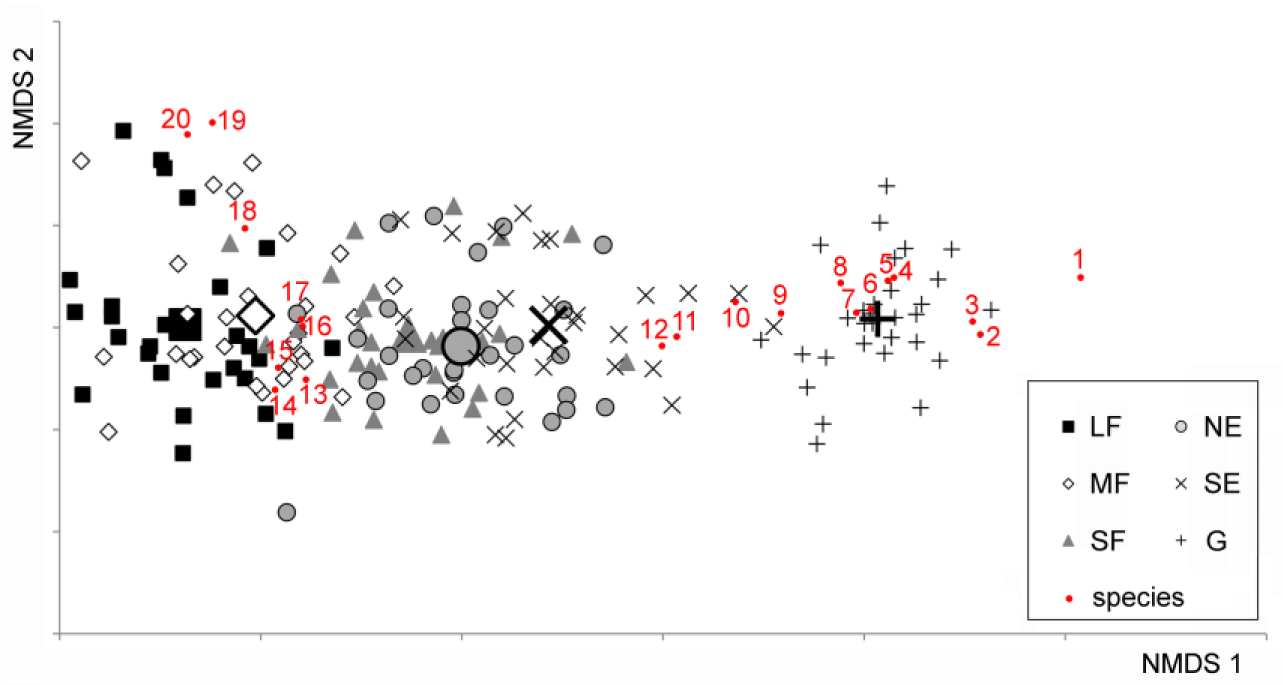
NMDS ordination scatterplot of the 176 plots. Only the top 20 species, according to the correlation to the ordination space (i.e., square root of the goodness of fit), are displayed. LF: large forest patches, MF: medium forest patches, SF: small forest patches, NE: north-facing edges, SE: south-facing edges, G: grasslands. The centroids for each habitat are drawn with larger signs. Species are as follows: 1: *Kochia laniflora*, 2: *Erophila verna*, 3: *Alkanna tinctoria*, 4: *Crepis rhoeadifolia*, 5: *Polygonum arenarium*, 6: *Holosteum umbellatum*, 7: *Poa bulbosa*, 8: *Syrenia cana*, 9: *Arenaria serpyllifolia*, 10: *Euphorbia seguieriana*, 11: *Stipa borysthenica* + *capillata*, 12: *Festuca vaginata*, 13: *Ligustrum vulgare*, 14: *Berberis vulgaris*, 15: *Rhamnus catharticus*, 16: *Celtis occidentalis*, 17: *Crataegus monogyna*, 18: *Bromus sterilis*, 19: *Stellaria media*, 20: *Anthriscus cerefolium*.

A considerable number of species were shared among all habitats (Figure 2). Woody habitats (i.e., forest patches plus forest edges) were strongly related, as shown by the high number of common species. The number of species restricted to single habitat types was also numerous, especially for north-facing edges, although grasslands, small forest patches and south-facing edges also had considerable numbers of species that did not occur elsewhere.

**Figure 2.**
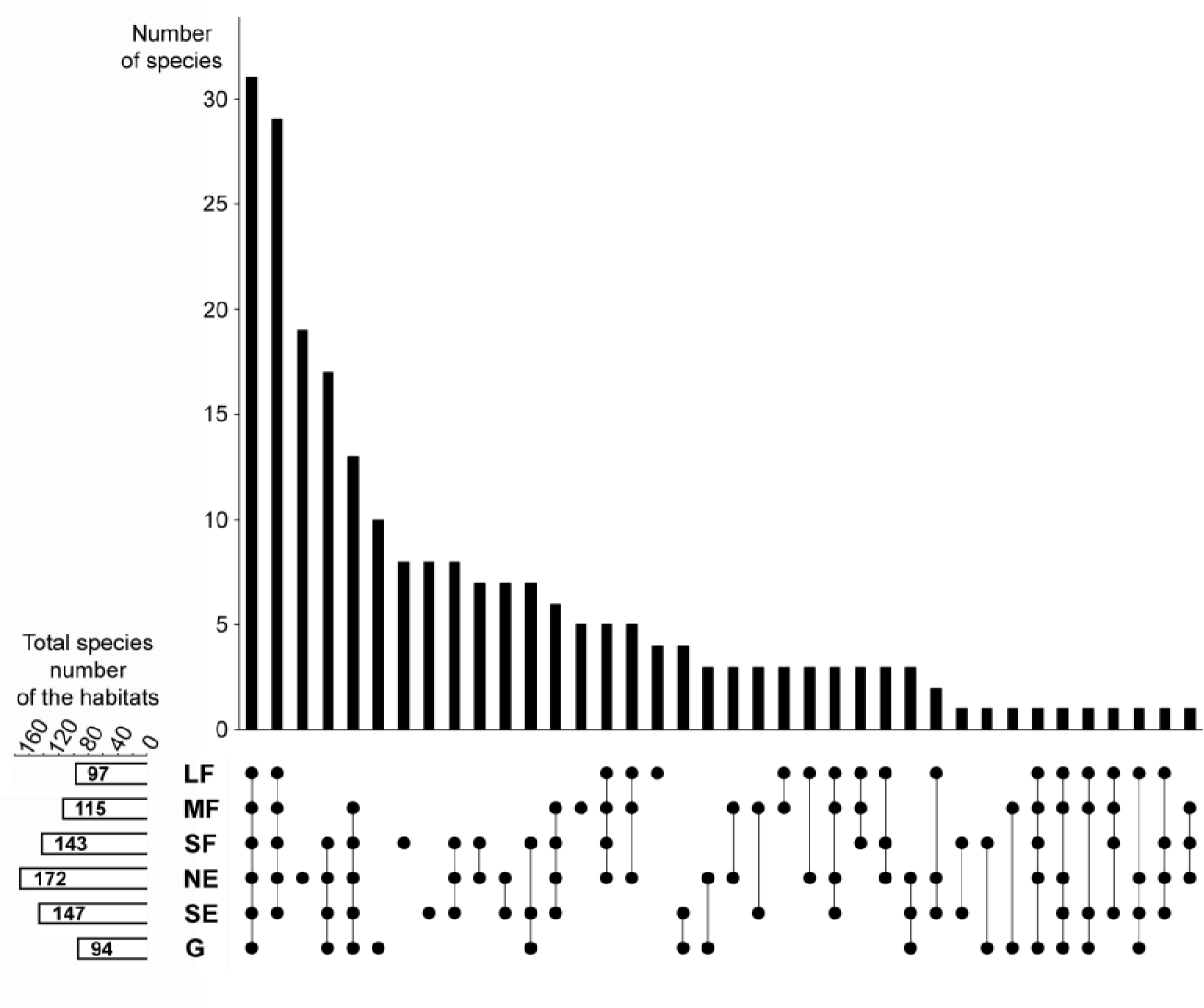
Relationships among the studied habitats in terms of species composition overlaps and distinctness. The upper panel shows the number of those species that were found in the habitats indicated by the dots in the lower panel. For example, the first column shows that there are 31 species that were found in all six habitats, whereas the third column shows that there are 19 species which are restricted to the north-facing grassland habitat. The small panel in the bottom left corner shows the total (i.e., pooled) number of species in each habitat. LF: large forest patches, MF: medium forest patches, SF: small forest patches, NE: north-facing edges, SE: south-facing edges, G: grasslands.

The total (i.e., pooled) number of species was the highest in north-facing edges (Figure 2), followed by south-facing edges and small forest patches. Medium forest patches had fewer species, while large forest patches and grasslands had the lowest total number of species.

Habitat type significantly influenced the mean ecological indicator values for temperature (F5,165 = 92.55, P < 0.001), soil moisture (F5,165 = 157.325, P < 0.001), and light availability (F5,165 = 226.47, P < 0.001). The mean ecological indicators for temperature showed that large and medium forest patches had the lowest, while grasslands had the highest values (Figure 3a). Small forest patches, north-facing forest edges and south-facing forest edges were intermediate, but their values were similar to those of the large and medium forest patches. The mean ecological indicators for soil moisture showed a continuously decreasing trend from the large and medium forest patches to the grassland, which had the lowest values (Figure 3b). Regarding the mean ecological indicators for light availability, there was a well-defined, gradually increasing trend from the large forest patches towards grasslands (Figure 3c).

**Figure 3.**
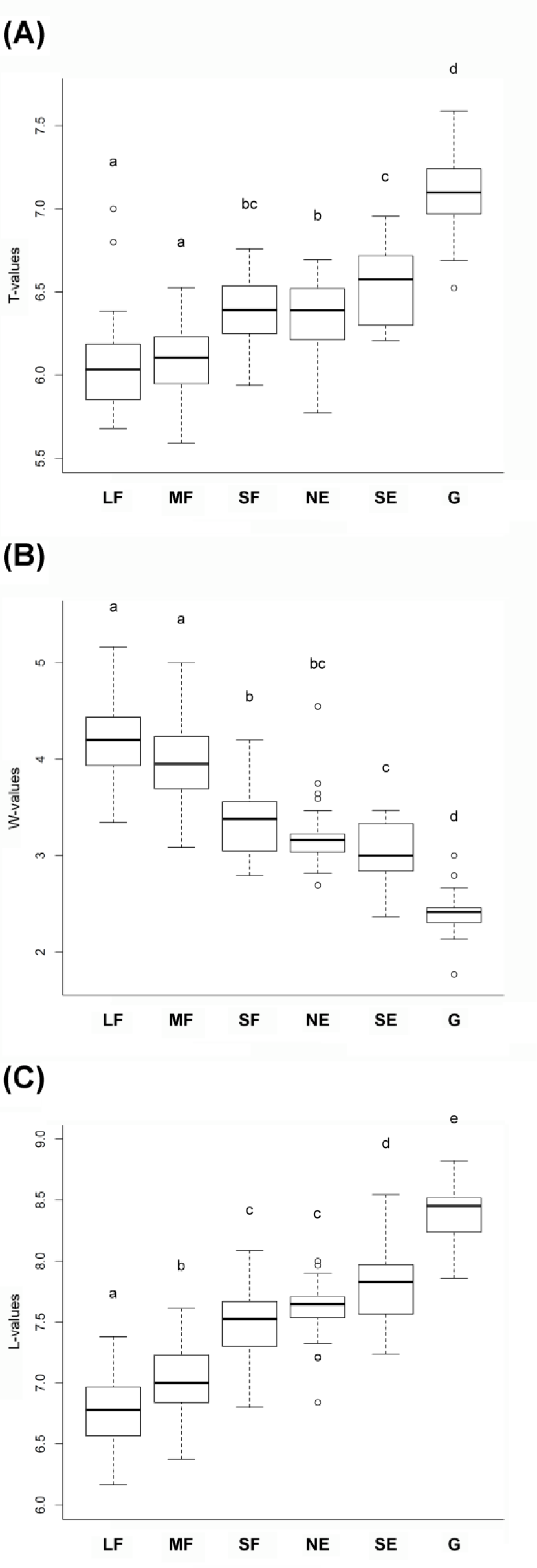
Mean ecological indicator values for (A) temperature, (B) soil moisture, and (C) light availability for the studied habitats. The habitats not sharing a letter are significantly different. LF: large forest patches, MF: medium forest patches, SF: small forest patches, NE: north-facing edges, SE: south-facing edges, G: grasslands.

Pearson’s chi-squared test showed that the frequency distributions of the life-form categories, geoelement types, and phytosociological preference groups in the six habitat types were different (chi-squared = 635.2, df = 30, P < 0.001; chi-squared = 339.2, df = 25, P < 0.001; chi-squared = 587.8, df = 25, P < 0.001, respectively). There was a continuous change in the life-form categories from the large forest patches towards the grasslands (Figure 4a). Large and medium forest patches did not differ significantly from each other, nor did small forest patches from north-facing edges. The proportions of shrubs and trees decreased towards the grasslands. Therophytes were most frequent in grasslands, while hemicryptophytes reached the maximum value in north-facing edges.

**Figure 4.**
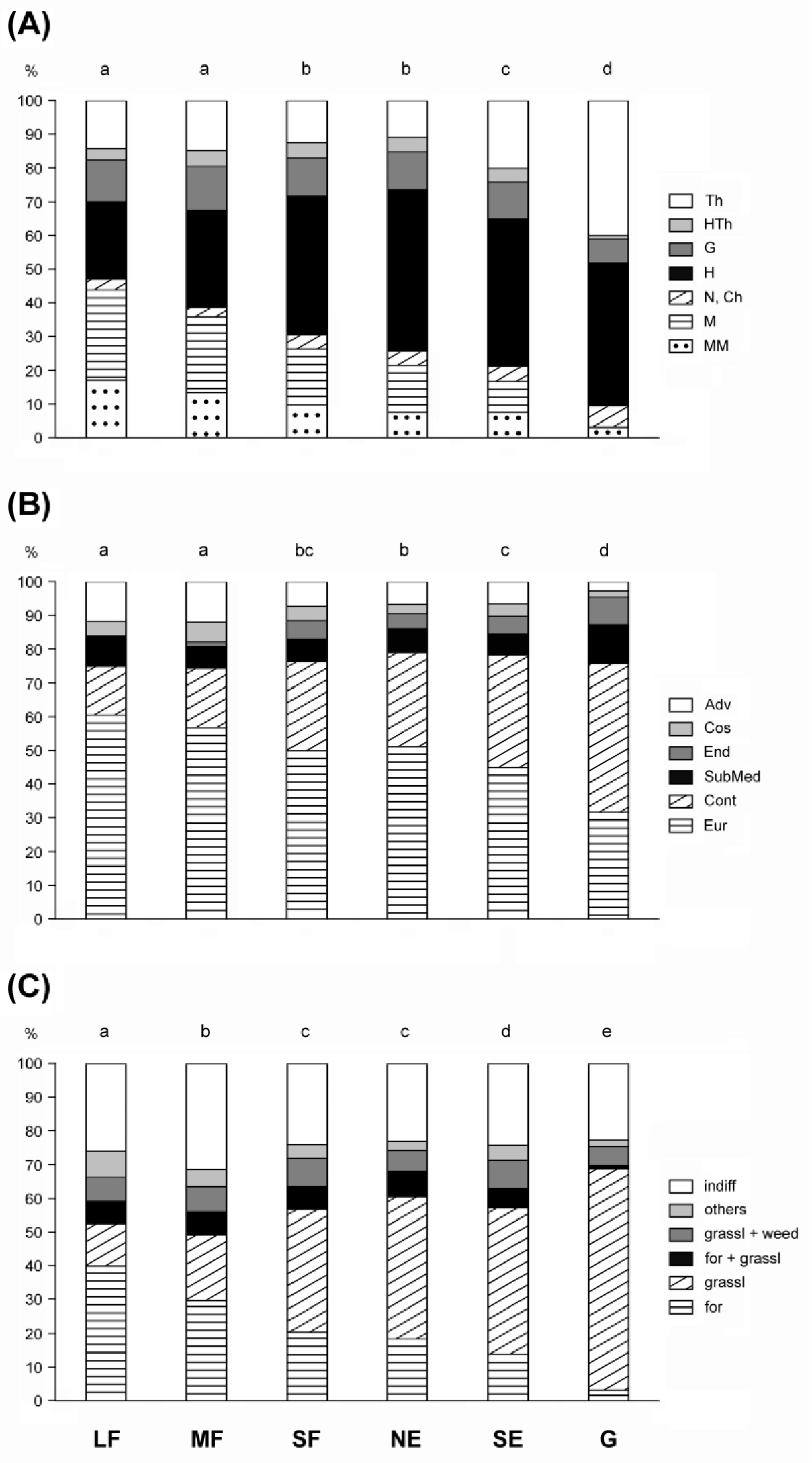
Frequency distributions for (A) life-form categories, (B) geoelement types, and (C) phytosociological preference groups of the six studied habitats. Th: therophytes, HTh: hemitherophytes, G: geophytes, H: hemicryptophytes, N: nanophanerophytes, Ch: chamaephytes, M: microphanerophytes, MM: meso- and megaphanerophytes; Adv: adventives, Cos: cosmopolitans, End: endemics, SubMed: submediterranean species, Cont: continental species, Eur: European and Eurasian species; indiff: indifferent species, others: species of edges, scrubs, and disturbed communities, grassl+weed: species related to grasslands and weed communities, for+grassl: species commonly found in forests and grasslands, grassl: grassland-related species, for: forest-related species; LF: large forest patches, MF: medium forest patches, SF: small forest patches, NE: north-facing edges, SE: south-facing edges, G: grasslands. The habitats not sharing a letter over the bars are significantly different.

The large forest patches were dominated by European and Eurasian species, the proportion of which became progressively smaller towards the grasslands (Figure 4b). In contrast, the proportion of continental species showed a reverse pattern. Other geoelements generally had a subordinate role. The grasslands contained the highest number of endemics and the lowest number of adventives.

The proportion of forest-related species showed a decreasing trend from the large forest patches towards the grasslands (Figure 4c). Species related to grasslands showed an opposite trend. Indifferent species (i.e., species with preferences towards multiple plant communities) played an important role in all studied habitats, but their proportion was the highest in medium forest patches.

## 4 Discussion

The components of forest-grassland mosaic ecosystems are conveniently classified into two basic categories: forest and grassland. The categorization is based on the dominant life-forms, but it correlates with numerous other characteristics, both biotic and abiotic (e.g. Breshears, 2006; Innes et al., 2013; Luza et al., 2014). While this categorization may be acceptable for some purposes, recent evidence has suggested that forest-steppes should not be regarded as simple two-phase systems (Erdős et al., 2018b).

Our study revealed a clear gradient regarding the studied characteristics. There was a mostly gradual species turnover from large forest patches to grasslands and gradual changes regarding ecological indicator values and functional species groups. Edges proved to be transitional between the forests and the grasslands, with south-facing edges being more similar to grasslands. Small forest patches represented a transition between medium forest patches and edges. Total species number peaked near the middle of the gradient, i.e., in north-facing edges.

In concordance with the mean ecological indicator values calculated in our study, earlier studies have also shown that forest interiors have lower temperature, and higher soil moisture than the neighboring treeless areas (Davies-Colley et al., 2000; Mosquera et al., 2014; Schmidt et al., 2017), whereas edges are generally intermediate regarding the above parameters (Kapos, 1989; Gehlhausen et al., 2000; Mosquera et al., 2014; Schmidt et al., 2017). In addition, we found that the indicator values of north-facing edges resembled those of the forest patches, which is in line with the findings of Matlack (1993). In contrast, the indicator values of south-facing edges were the most similar to those of the grasslands. Ries et al. (2004) predicted that south-facing edges should deviate from forest interiors more than north-facing ones, due to the larger exposure to sunlight Thus it seems likely that the studied habitats represent a continuous gradient of environmental factors, ranging from the largest forest patches to the open grasslands. This may have extremely important consequences. Forest-steppes have very high species richness (Zlotin, 2002; Erdős et al., 2018a) and habitat heterogeneity has been suggested as an important driver of this high species diversity (Tölgyesi et al., 2018; Erdős et al., 2018b). Our results reinforce and complement these findings, emphasizing that a large number of species can find appropriate habitats somewhere along the gradient, provided that the full range of heterogeneity is preserved in the mosaic.

For most of the studied characteristics, small forest patches were similar to north-facing edges. This result indicates that small forest patches are edge-like habitats; that is, they are too small to have a core area. It has been shown that the microclimate of the outer areas of forest patches is strongly influenced by a width of dozens of meters (Ries et al., 2004; Hennenberg et al., 2008; Dodonov et al., 2013; Schmidt et al., 2017).

As shown by the geoelement spectra, the sandy forests in our study region are dominated by European and Eurasian species, while the sandy grasslands are richer in continental species. This phenomenon can be explained by biogeographic causes. Deciduous forests extend as a wide belt from Western Europe to Inner Asia (Walter and Breckle, 1989; Schultz, 2005). In contrast, natural grasslands occur from eastern Europe to the Far East, while most Central and Western European grasslands are secondary and depend on human activities such as grazing and mowing. The grasslands in the Carpathian Basin may be understood as the westernmost extensions of the continental grasslands.

The sequence of habitats revealed in this study may be termed a coenocline (Whittaker, 1967, 1975). The existence of a quite continuous gradient of the studied habitats suggests that the dichotomic categorization of forest-steppe habitats into forest and grassland patches is a serious and misleading oversimplification, as it is a poor representation of the real heterogeneity of the forest-steppe ecosystem. We believe that forest-steppe patterns better fit the gradient-based paradigm of landscape structure (Cushman et al., 2005; McGarigal and Cushman, 2005), which, rather than using dichotomic categorizations, is able to reflect more continuous variations. More specifically, by using the forest vs. grassland categories, all forest patches (irrespective of their sizes), and the edges are classified into the forest category, disregarding the variability among them, which results in a loss of information (Cushman et al., 2005). Thus, looking beyond the forest-grassland dichotomy seems necessary for a better and more correct understanding of forest-steppe ecosystems. Our study indicates that the gradient-based paradigm may prove useful in other forest-grassland mosaic ecosystems such as savannas, wood pastures, or prairie-forest ecotones. Similarly, the gradient-based approach can be of importance in forest ecosystems suffering from heavy anthropogenic fragmentation.

Regarding the vegetation gradient revealed in this study, the question emerges as to what the drivers of this gradient are. In the sand dune areas of the Kiskunság region, background factors such as groundwater depth, soil moisture content, and microclimatic parameters strongly depend on terrain features (e.g., Bátori et al., 2014; Tölgyesi et al., 2014). However, both our field experience and earlier research (e.g., Halupa, 1967; Bölöni et al., 2011) show that patches of poplar forests can be found in various topographical positions: on sand dune tops and ridges, on windward and leeward dune slopes, and in dune slacks, even though these positions differ strongly in terms of abiotic parameters. In other words, there is no apparent relation between the presence of differently sized forest patches and current environmental parameters. The probable explanation for this is twofold. First, the horizontal roots of *Populus alba* may extend up to 40 m or more (Magyar, 1961; Halupa, 1967; Szodfridt, 1969). Thus, trees situated in a hostile environment (such as a dry dune ridge) are able to reach soils with a higher humus content and better moisture supply (for example, in a dune slack). Second, forest patches sometimes depend on one or more humus layers buried under the sand (Szodfridt, 1969; Halupa, 1967; Bodrogközy, 1982; Molnár, 2003). These buried layers originate from earlier vegetation periods, as the wind has re-deposited the sand dunes several times during the Holocene (Molnár, 2009; Molnár et al., 2012). In summary, the current pattern of differently sized forest patches may reflect an earlier sand dune topography and associated environmental parameters. Our study has focused on forest habitats, analysing only one grassland type, the open perennial sand grassland, which dominates the study sites. Other herbaceous habitats play a subordinate role in the study sites. It can be assumed that non-forest habitats also form a gradient. For example, *Salix rosmarinifolia* subshrub communities occur mainly in dune slacks with favorable water availability (Bodrogközy, 1982; Borhidi et al., 2012). Closed sand steppes thrive under semi-dry conditions (Borhidi et al., 2012) and have been shown to be compositionally related to forest edges (Erdős et al., 2013), while open perennial sand grasslands and annual sand grasslands live under the harshest circumstances. Also, there may be considerable differences between the differently sized grasslands, with the smallest grassland patches of natural openings resembling forest edges (Molnár, 1998). The possible gradient-like arrangement of non-forest communities in the studied mosaic ecosystem calls for further studies to understand forest-steppe heterogeneity in the frame of the gradient-based paradigm of landscape structure.

## 6 Author Contributions

LE, GK-D, and PT conceived the research idea. LE, GK-D, ZB, CT, PT, and PJK collected the data. KS, ÁB-F, CT, and PJK performed the statistical analyses. LE and PT, with contributions from GK-D, wrote the paper. All authors discussed the results and commented on the manuscript.

## 8 Acknowledgments

The authors thank Dolly Tolnay and Mihály Szőke-Tóth for their help with the field work, and Attila Lengyel (MTA Centre for Ecological Research) for his remarks concerning the ordination.

